# Molecular pet or parasite? Exploring selection for vertical and horizontal plasmid transfer

**DOI:** 10.64898/2026.06.02.729688

**Authors:** Elizabeth S. Duan, Eva M. Top, Benjamin Kerr, Olivia Kosterlitz

## Abstract

Understanding the environmental conditions that drive selection for increased horizontal plasmid transfer is crucial for predicting the spread of plasmid-encoded antibiotic resistance. In natural systems, plasmids exhibit diverse lifestyles, ranging from “host-centric” strategies, which favor vertical gene transfer (VGT) from mother to daughter cell at the expense of horizontal mobility, to “parasitic” strategies, which favor horizontal gene transfer (HGT) by conjugation at the expense of host fitness. However, laboratory evolution experiments are biased towards host-centric evolution, highlighting a gap in our ability to consistently select for horizontal mobility. To understand this experimental bias, we developed a mathematical model to explore the invasion of pleiotropic transfer mutations. Using local linear stability analysis, we derived an invasion criterion establishing that, for a given pleiotropic cost, the availability of plasmid-free cells determines whether increased transfer is selected. We expanded this model to better represent our previous evolution experiment, in which selection for a host-centric mutant occurred despite the addition of plasmid-free cells and periodic selection for transconjugants. We found that standard batch culture protocols inherently impose strong selective pressure on VGT, heavily limiting the laboratory observation of increases in HGT. We experimentally and theoretically demonstrated that a simple protocol modification—minimizing excess growth by eliminating batch culture passages—effectively tips selection towards HGT. Finally, we performed a parameter sweep to predict the invasion success of hypothetical mutants across HGT-VGT phenotypic space. Our predictive framework can be used to further explore the evolution of plasmid transfer under conditions more representative of natural environments where medically and environmentally relevant plasmids evolve.

## Introduction

A defining feature of conjugative plasmids is that these extrachromosomal elements encode machinery for their own horizontal gene transfer (HGT). Because such plasmids are primary vehicles for interspecies spread of antibiotic resistance, understanding the evolutionary processes that influence HGT rates is essential for public health [1,2]. HGT rates have been observed to vary over 13 orders of magnitude, suggesting that some natural environments select for higher HGT rates [3,4]. However, experimental plasmid evolution in laboratory conditions often results in reduced HGT rates, indicating a gap in our understanding of how to place strong selective pressure on HGT. Understanding how environmental conditions, whether natural or artificial, favor higher or lower HGT rates will clarify the selective forces shaping plasmid evolution and help explain why laboratory evolution often fails to reproduce the high transfer rates observed in nature.

One factor that may result in reduced HGT rates in laboratory evolution experiments could be the potential fitness costs associated with HGT. HGT requires the translation of plasmid-encoded conjugation machinery, including proteins dedicated to pilus assembly, membrane translocation, and transfer operon regulation [5]. Host resources are used to build this machinery, which could take away from the host’s ability to grow [4,6]. Many evolution experiments employ serial transfer in liquid batch culture conditions [7], which places strong selection for host growth and thus could select against costly HGT. As plasmids are also inherited during host cell division (vertical gene transfer, VGT), it may be more beneficial for plasmids to reduce their HGT rates for increased VGT in these standard laboratory conditions. This “host-centric” evolution has been observed through compensatory evolution (mutations that reduces plasmid fitness cost) in previous experiments [8–13]. This indicates potential pleiotropy between the two traits—a change in VGT affecting HGT.

We can visualize pleiotropic mutations as points in a two-dimensional phenotypic space, where the axes correspond to horizontal and vertical plasmid transfer (Figure 1). Setting the ancestor as the origin, mutations that increase at least one mode of transfer would land in quadrants (Q) 1, 2, or 4. Improving both modes of transfer indicates a “synergistic” evolutionary outcome, where the overall phenotypic movement due to one or more mutations places descendants in Q1. Alternatively, cases of antagonistic pleiotropy indicate an increase in one mode of transfer that comes at the expense of the other. In this paper, we term evolutionary outcomes that increase VGT at the expense of HGT “host-centric” (Q2), while those that increase HGT at the expense of VGT are termed “parasitic” (Q4). Synergistic outcomes would be favored in any environment, but they are rarely observed compared to antagonist outcomes, potentially due to a tradeoff between HGT and VGT. Antagonistic outcomes are also more informative, as increasing one mode of transfer at the expense of the other indicates the environment selects strongly for the favored mode of transfer. Thus, we focus on host-centric and parasitic outcomes to better understand how environmental conditions place selection on VGT or HGT. While selective conditions favoring host-centric evolution are well understood, defining the precise conditions that drive parasitic evolution is still a challenge.

**Figure 1.**
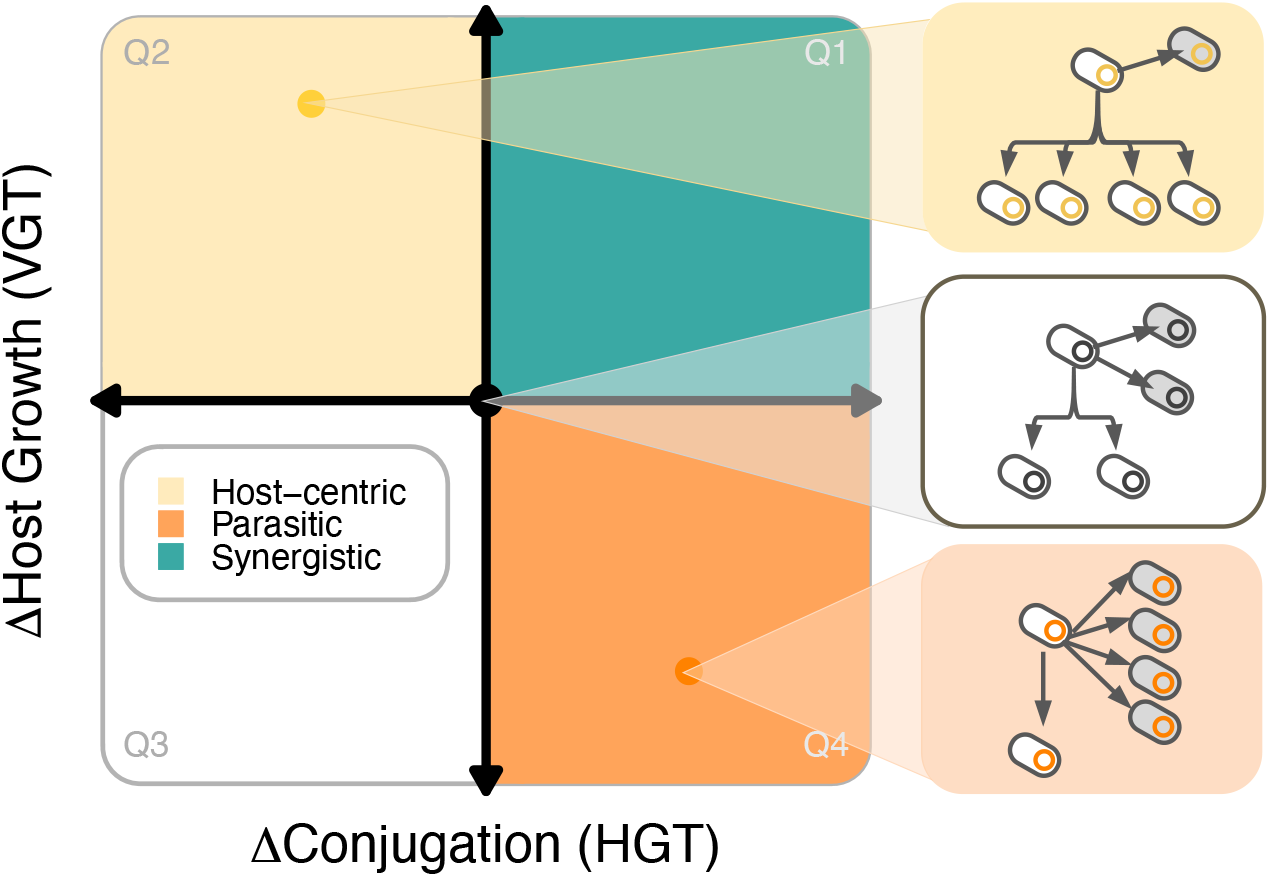
The effects of pleiotropic transfer mutations can be represented as points in a two-dimensional plane. Axes can be centered at the conjugation and host growth rates of an ancestral strain, with the x-axis representing changes in conjugation rate (HGT) and the y-axis representing changes in host growth rate (VGT). Mutations that land in quadrant 1 (teal) are synergistic, increasing both HGT and VGT, and thus are always predicted to be favored. Mutations in quadrant 3 (white) decrease both and are never expected to be favored. We focus on mutations that exhibit antagonistic pleiotropy to determine whether environmental conditions place stronger selection on VGT or HGT. When conditions favor VGT, host-centric evolution that increases VGT but decreases HGT can occur (Q2, yellow quadrant). When conditions favor HGT, parasitic evolution that increases HGT but decreases VGT can occur (Q4, orange quadrant). In the insets, vertical arrows connect mother to daughter cells while horizontal arrows indicate conjugation into neighboring cell(s); the number of arrows relates to the rate of VGT or HGT.

To identify conditions that drive selection towards HGT, we consider HGT-proficient plasmids to be like parasitic agents. Parasite evolution is often driven by the availability of susceptible hosts [6,14,15]. For conjugative plasmids, plasmid-free cells can be likened to susceptible hosts. Thus, including plasmid-free hosts in evolution experiments can offer the opportunity to transfer horizontally and allow parasitic advantage to be realized [6,16]. Some protocols have additionally implemented forced conjugation by applying antibiotic selection for plasmid-free cells that received the plasmid (i.e., transconjugants), which makes HGT a requirement for plasmids to escape extinction during the experiment [17,18]. While implementing these conditions has led in some cases to selection for parasitic mutations, some studies still observe host-centric evolution where conjugation decreases [6,17,19]. This difficulty in consistently selecting for HGT highlights a gap in our understanding of selection for increased horizontal transfer.

In this study, we aim to further our understanding of selection for HGT. We begin by developing an abstract mathematical model to derive a general selection criterion for pleiotropic transfer mutations. We apply these findings in a case study using strains from our previous plasmid evolution experiment conducted by De Gelder et al. [17]. As these strains differ by a single mutation and represent different plasmid lifestyles relative to each other (parasitic vs host-centric), we can analyze their competitive success under various conditions to understand how these conditions impose selective pressure on HGT or VGT. Finally, we expand our case study to include hypothetical mutations spanning phenotypic space, providing a broader perspective on how environmental conditions dictate horizontal transfer selection.

## Results

### A simplified mathematical model to predict mutant plasmid invasion

To identify general factors favoring selection for vertical versus horizontal transfer, we start with a simple and abstract mathematical model in which plasmid mutations can simultaneously affect both VGT and HGT. In this model, we analytically explore the conditions under which a plasmid with a mutation that alters the rates of conjugation (HGT) and host growth (VGT) could invade a population of cells possessing the ancestral plasmid.

We start by tracking three bacterial populations: cells with the ancestral plasmid (density *A*), cells with the mutant plasmid (density *M*), and cells with no plasmid (density *N*). We use lower-case letters to represent the proportions of each of the three strains in the community:

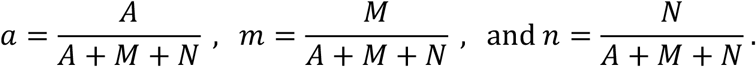

We label our three bacterial strains with non-italicized letters A, M, and N. Letting I and J denote arbitrary strains, we can characterize key parameters. We let *ψ*_I_ and *δ*_I_ be the growth and death rates of strain I (with I ∈ {A, M, N}). We let *γ*_J_ be the rate that a plasmid transfers from a plasmid-bearing cell of strain J to a plasmid-free cell; and *σ*_J_ be the probability that, upon cell division, a parent cell of strain J generates an offspring cell with no plasmid (with J ∈ {A, M}).

The dynamics of our bacterial community are given by the following differential equations (Figure 2A):

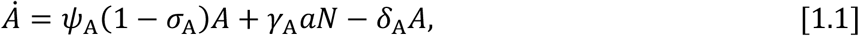

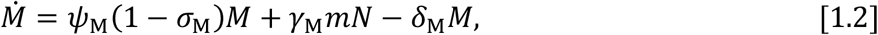

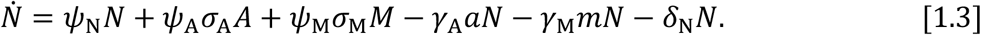

**Figure 2.**
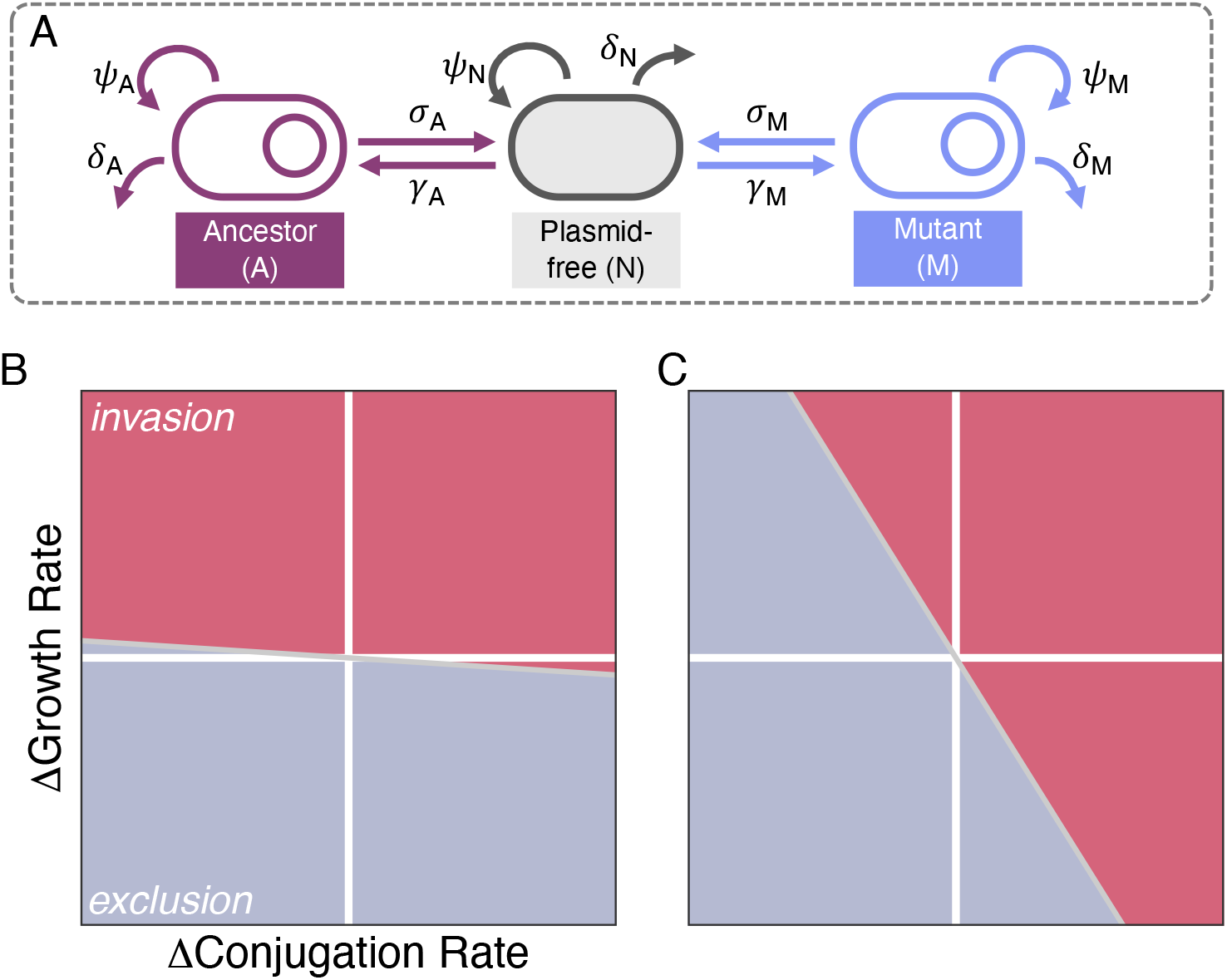
An abstract mathematical model allows us to derive an invasion criterion for pleiotropic transfer mutations. (A) Our system consists of ancestral plasmid-bearing (*A*), mutant plasmid-bearing *(M*), and plasmid-free (*N*) cells whose dynamics are defined by growth (*ψ*), conjugation (*γ*), segregation (*σ*), and death (*δ*). (B) Reducing access to plasmid-free cells decreases the magnitude of the slope of the invasion boundary (gray line, condition [4]), favoring host-centric and synergistic mutations. (C) Alternatively, increasing access to plasmid-free cells increases the magnitude of the slope, favoring parasitic over host-centric mutations. In both parts B and C, pleiotropic mutations in the red regions invade, while those in the blue regions fail to do so.

These equations make some simplifying assumptions about our populations. First, we assume population growth is exponential (i.e., without density dependence). Second, conjugation rate is assumed to depend on the proportion of the relevant plasmid-bearing cell (variables *a* and *m*), as opposed to its density (variables *A* and *M*). Although densities of donors and recipients influence the production of transconjugants under some conditions, there also may be circumstances where horizontal transmission is more frequency-than density-dependent [20]. These simplifications enable us to derive an analytical invasion criterion for pleiotropic transfer mutations.

### Defining an invasion criterion for pleiotropic transfer mutations

We perform a local linear stability analysis to determine when a rare pleiotropic transfer mutation is expected to invade. We first consider a community of only ancestral plasmid-bearing and plasmid-free cells and solve for the equilibrium in which both strains stably coexist. Then, we introduce a small number of mutant plasmid-bearing cells (representing de novo mutation) and determine the stability of the original equilibrium with mutant plasmid-bearing cells now present. If the original equilibrium is still stable, the mutant perturbation decays and the system returns to a community with only ancestral plasmid-bearing and plasmid-free types, representing mutant exclusion. If the original equilibrium is unstable, the mutant perturbation grows and the system departs from the original equilibrium, representing mutant invasion.

We begin by defining our focal mutation in terms of changes to ancestral conjugation and growth rates (i.e., *γ*_M_ = *γ*_A_ + Δ_*γ*_ and *ψ*_M_ = *ψ*_A_ + Δ_*ψ*_). For ease, we will also assume here *σ*_A_ = *σ*_M_ = *σ* and *δ*_A_ = *δ*_M_ = *δ*; that is, the mutation on the plasmid does not affect segregational loss nor the death rate of the host cell. Using all this information, we show in Supplemental section S1 that this model can be recast in terms of strain proportions:

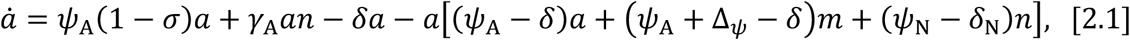

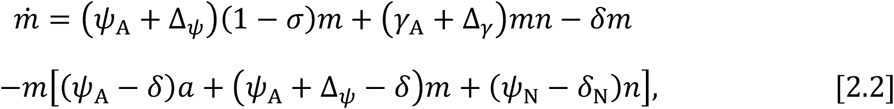

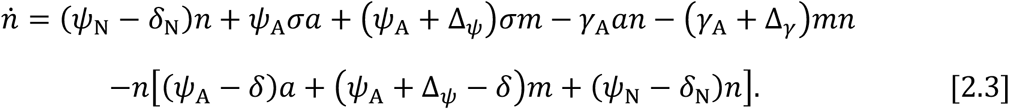

In a community without mutant plasmids present (*m* = 0), there are two possible equilibria for equations [2]. The first involves a population of only plasmid-free cells (i.e.,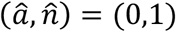). The second is the following non-trivial equilibrium (see Supplemental section S2):

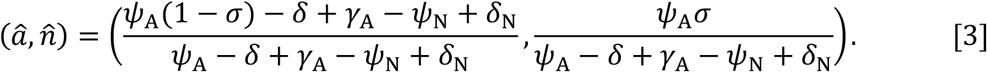

If the net growth rate of the ancestor outpaces its death (i.e., *ψ*_A_(1 − *σ*) > *δ*) and the segregation rate is positive (i.e., *σ* > 0), then it follows that equilibrium [3] involves coexistence of ancestral-plasmid-bearing and plasmid-free cells (i.e., 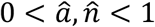). Without mutant plasmids present, this second equilibrium is stable (see Supplemental section S3).

We then explore what happens when a small proportion of mutant plasmid-bearing cells is introduced. A local linear stability analysis in Supplemental section S3 demonstrates that the condition for mutant invasion is:

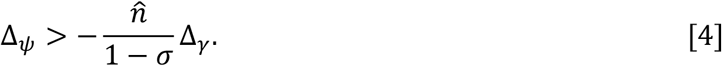

We explore the implications of this invasion criterion for various pleiotropic transfer mutations.

First, a synergistic mutation improves both modes of transfer (Δ_*γ*_ > 0 and Δ_*ψ*_ > 0). Such a mutation will always satisfy condition [4] and therefore will always invade. Results are less straightforward for mutations involving antagonistic pleiotropy. Thus, we next recast the condition [4] for parasitic and host-centric mutations to determine conditions that favor HGT at the expense of VGT, and those that favor VGT at the expense of HGT.

A parasitic mutation improves conjugation at a growth cost (Δ_*γ*_ > 0 and Δ_*ψ*_ < 0). In such a case, condition [4] can be reworked as:

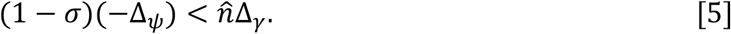

Condition [5] states that the growth cost of the mutation (the left-hand-side of [5]) must be less than the conjugation benefit (the right-hand-side of [5]). The benefit is weighted by the proportion of plasmid-free cells, as the effect of the mutation is realized by converting plasmid-free cells at a greater rate. The cost of the mutation is weighted by one minus the segregation probability, which is the fraction of the growing population that remains plasmid-bearing. Because 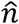 increases as *ψ*_N_ – *δ*_*N*_ increases, the invasion criterion is easier to satisfy as the plasmid-free cell population increases its net growth, either via increased growth or decreased death. Lastly, as segregation loss increases (as *σ* goes up), the criterion becomes easier to satisfy for two reasons. First, 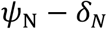 increases as *σ* increases, so the right side grows as *σ* increases. Second, the left side shrinks as *σ* increases (the growth penalty of the mutant plasmid is more muted when plasmid-bearing cells are lost through division at a greater rate).

Finally, a host-centric mutation improves host growth at the expense of conjugation (Δ_*ψ*_ > 0 and Δ_*γ*_ < 0). In this case condition [4] can be written as follows:

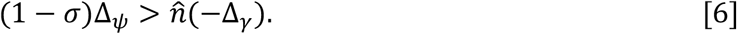

Condition [6] simply reverses the logic of condition [5]: now the growth benefit needs to outweigh the cost to conjugation. All the same factors (e.g., plasmid-free net growth, segregational loss) impact whether condition [6] holds, where the changes in parameters that favor parasitic mutations now have the opposite effect on host-centric mutations.

Overall, the amount of plasmid-free cells available as recipients for conjugation impacts the kind of pleiotropic mutation that is favored. Specifically, as 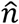 in [4] decreases (e.g., due to low plasmid-free net growth or low segregational loss), the negative slope of the boundary giving mutant invasion becomes increasingly shallow. As shown in Figure 2B, this favors mutations that improve growth (even if at the expense of transfer). On the other hand, as 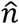 increases (e.g., due to high plasmid-free net growth or high segregational loss), the boundary becomes steeper and mutations that improve transfer are favored (even if at the expense of host growth; see Figure 2C). Thus, this model highlights a clear prediction with regards to the role of available plasmid-free hosts on the evolutionary trajectory of pleiotropic mutations in the plasmid.

In this abstract model, plasmid-free cell availability is primarily governed by their net growth rate and segregational loss of the plasmid. However, in laboratory experiments, plasmid-free cells are often made available by exogenous addition. Does such a change alter the basic expectations of our abstract model? To explore this question, we focus first on an experiment with such exogenous addition as a “case study.” This case study motivates a more biologically accurate model to discern how both the nature of pleiotropy and experimental conditions impact selection for different modes of plasmid transfer.

### Case study on strains from our previous plasmid evolution experiment

To compare our abstract model findings to an experimental system, we turned to our previous evolution experiment, De Gelder et al. [17], which evolved the IncP-1β plasmid pB10 in *Stenotrophomonas maltophilia* P21. The study aimed to understand how host-switching influences the evolution of long-term host range and thus implemented a multi-phase protocol that included both daily batch culture passages and forced conjugation through transconjugant selection to move evolving plasmids into new hosts (Figure 3A). As a result, the protocol simultaneously imposed selection for vertical transfer (during batch culture growth) and horizontal transfer (through the introduction of plasmid-free hosts and strong transconjugant selection with donor counterselection).

**Figure 3.**
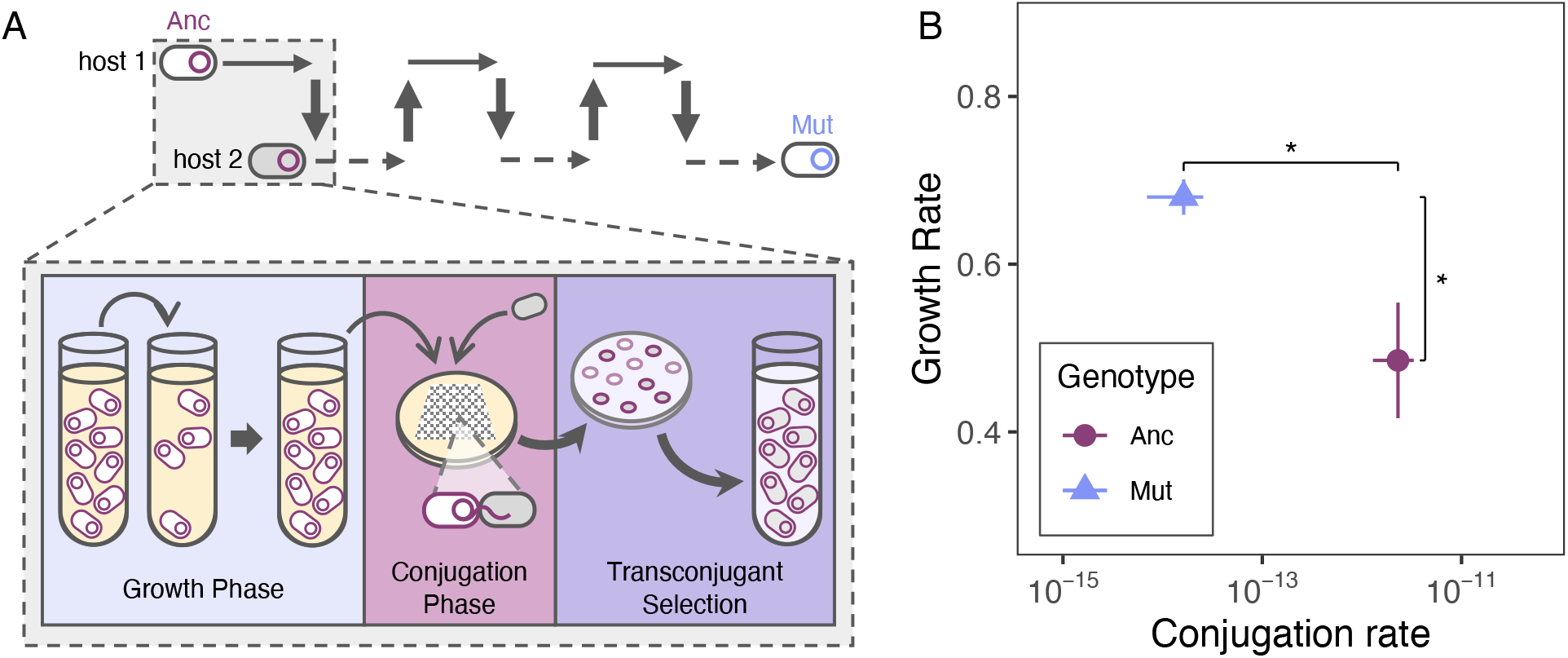
A previous evolution experiment by De Gelder et al. (2008) selected for a host-centric mutation. (A) The ancestral pB10 (Anc, red) in *S. maltophilia* was evolved in a protocol cycle with host-switching (figure adapted from De Gelder et al.) Specifically, the protocol included growth (light blue background), conjugation (light pink background), and transconjugant selection (purple background) phases. The resulting mutant (Mut, blue) was selected for. (B) The mutant has a significantly higher growth rate and significantly lower conjugation rate compared to the ancestor, indicating host-centric evolution. Error bars represent standard error of the mean.

We measured the growth and conjugation rates of evolved and ancestral plasmids in the ancestral host and found that one mutant decreased its conjugation rate (from 2.3 x 10^-12^ to 1.63 x 10^-14^ mL CFU^-1^ hr^-1^, *p* = 0.03, Welch’s t-test) while increasing its growth rate (from 0.48 to 0.62 hr^-1^, *p* = 0.02, Welch’s t-test), displaying host-centric evolution (Figure 3B). This ancestral-mutant plasmid pair differs by one mutation (A25E in the *trbC* prepilin gene). The success of this host-centric mutant—a better grower but worse conjugator—indicates that despite explicit selection for HGT via forced conjugation, the De Gelder et al. protocol still applied greater selection for VGT than HGT.

### Long growth phases drive selection for VGT

To understand why the host-centric mutant was selected for in the De Gelder et al. protocol, we simulated the invasion of the mutant using an expanded version of our abstract model that is more representative of experimental conditions. We tracked the densities of three bacterial populations: cells bearing the ancestral plasmid, cells bearing the mutant plasmid, and plasmid-free cells (denoted A, M, and N, respectively). As before, we will let I and J be generic variables for these strains (e.g., I ∈ {A, M, N}). The concentration of a resource is given by a dynamical variable *C*. Growth rate for each strain decreases with resource concentration according to a Monod equation:

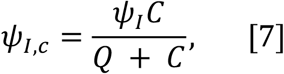

Where *ψ*_I_ is the maximum rate of growth for strain I and *Q* is the half-saturation constant (assumed to be the same for all strains). As *C* approaches zero, growth stops, representing stationary phase. We represented batch culture transfer by including specifications for dilution rate and time frames between dilutions: after a specified amount of time, a dilution is performed and resources are replenished. Additionally, we specified two host cell types (1 and 2) to represent alternating donors and transconjugants; we add this host-cell type as a subscript on our dynamic variables, *A, M*, and *N*. Using the same parameter designations as our abstract model above, the updated differential equations are as follows:

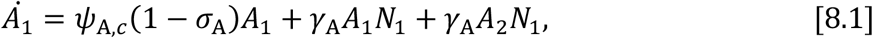

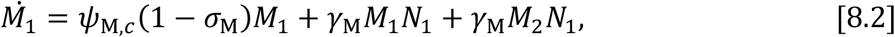

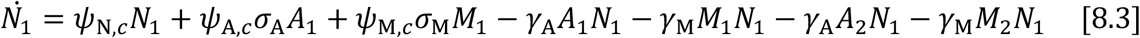

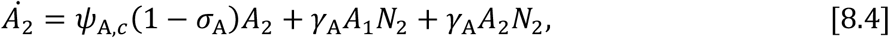

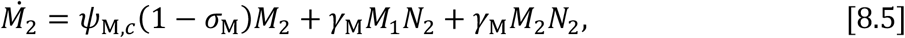

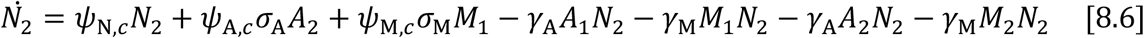

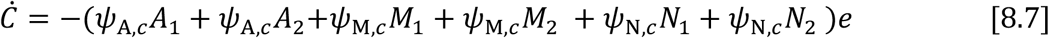

The parameter *e* is the specific consumption rate (i.e., inverse of the yield coefficient). We programmed each phase of the De Gelder et al. protocol into our simulations (see Materials and Methods). For our experimental simulations, we assume segregational loss is not present (*σ*_I_ = 0 for I ∈ {A, M}). We also assume that the addition of transconjugant-selecting antibiotics immediately kills all donors and recipients. Thus, plasmid-free cells are only present due to exogenous addition and available for conjugation during conjugation phases.

The De Gelder et al. protocol consisted of a growth phase, conjugation phase, and transconjugant selection phase (Figure 3A). We found that the mutant increased in frequency during growth phases but decreased in frequency during conjugation phases (Figure 4). Due to the length of the growth phase (seven rounds of serial passages), the host-centric mutant was able to invade and outcompete the ancestor.

**Figure 4.**
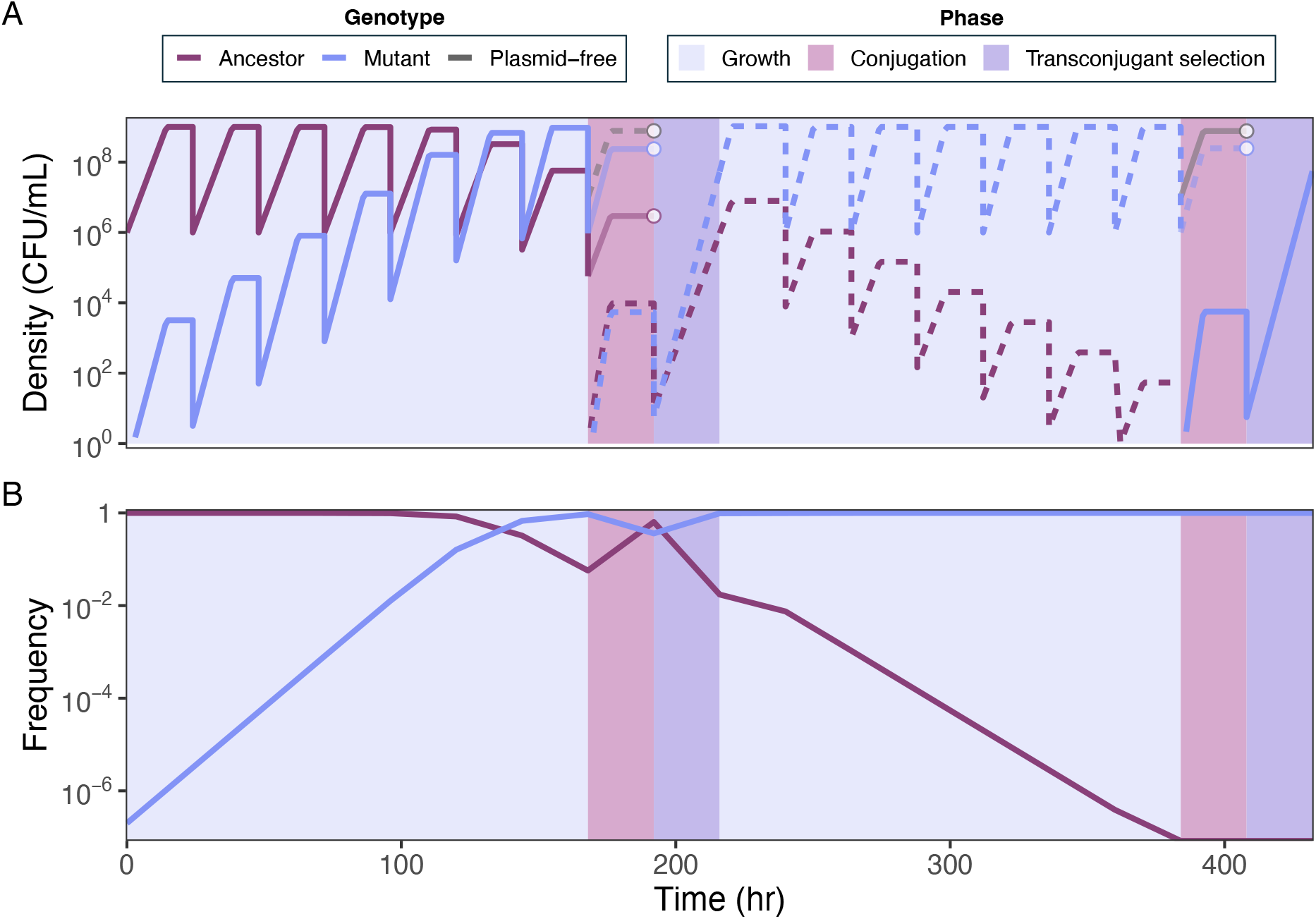
The host-centric mutant invades in De Gelder et al. due to long growth phases. (A) Simulated cell densities in two cycles of the evolution experiment. The simulation begins with one cell of the mutant (better grower, blue) in a large population of ancestor cells (better conjugator, red). In growth phases (light blue background), strains simply grow until resources are exhausted. In conjugation phases (pink background), plasmid-free cells (gray) of the opposite cell type (alternating dashed and solid) are added to allow conjugation to occur. Donors and recipients to be selected against in the following transconjugant selection phase (purple background) have lightened densities during conjugation phases, and the open point represents their immediate death upon entry to the transconjugant selection phase. (B) Frequencies of plasmid-bearing types calculated from densities. Frequencies are plotted every 24 hours. In conjugation phases, the frequency of the selected transconjugant type is plotted at the end of the phase.

### Removal of growth phases shifts selection to favor HGT

Even with the periodic addition of plasmid-free cells and forced conjugation, the De Gelder et al. protocol did not select for HGT over VGT. Our simulations show that the extended growth phase contributed strongly to selection of the host-centric mutant. Thus, we hypothesized that removing the growth phase entirely would decrease emphasis on growth and shift selection to favor HGT over VGT.

We modified the De Gelder et al. protocol by removing all growth phases. The De Gelder et al. experiment represents a “Low Frequency Conjugation” (LFC) protocol, with a growth phase (seven rounds of serial passage) before one conjugation phase and transconjugant selection phase. Our new “High Frequency Conjugation” protocol, with no growth phase, is expected to favor HGT by decreasing the emphasis on growth and increasing the frequency of conjugation and transconjugant selection phases.

To confirm the selective pressure of each protocol, we ran invasion experiments. These experiments are set up with a resident strain and an invading strain. The resident strain is numerically dominant, comparable to the ancestral strain in an evolution experiment. The invading strain is rare compared to the resident, and comparable to a mutant that arose. We compete the resident and invading strains in a protocol predicted to favor the invading strain and assess whether the invading strain increases in frequency (successful invasion). For the LFC, we expect selection to favor VGT, so a host-centric invader is predicted to outcompete the resident strain. Oppositely, in the HFC, we expect to favor HGT, so a parasitic invader is predicted to outcompete the resident strain.

As the ancestor and mutant strains from De Gelder et al. exhibit antagonistic pleiotropy relative to each other, we can use these strains in various roles to test the selective pressures of the protocols. Moving forward, we refer to the ancestor as genotype X (the better conjugator but worse grower, named “X” because it has the higher value on the *x*-axis), and the mutant as genotype Y (the better grower but worse conjugator, named “Y” because it has the higher value on the *y*-axis). We use these abstract symbols because their roles (resident or invader) will change in each experiment.

For the LFC invasion experiment, the resident strain is represented by genotype X, which places the invading genotype Y in the host-centric quadrant with respect to X (Figure 5A). For the HFC experiment, we flip the strain roles such that genotype Y is the resident strain, placing the invading genotype X in the parasitic quadrant (Figure 5C).

**Figure 5.**
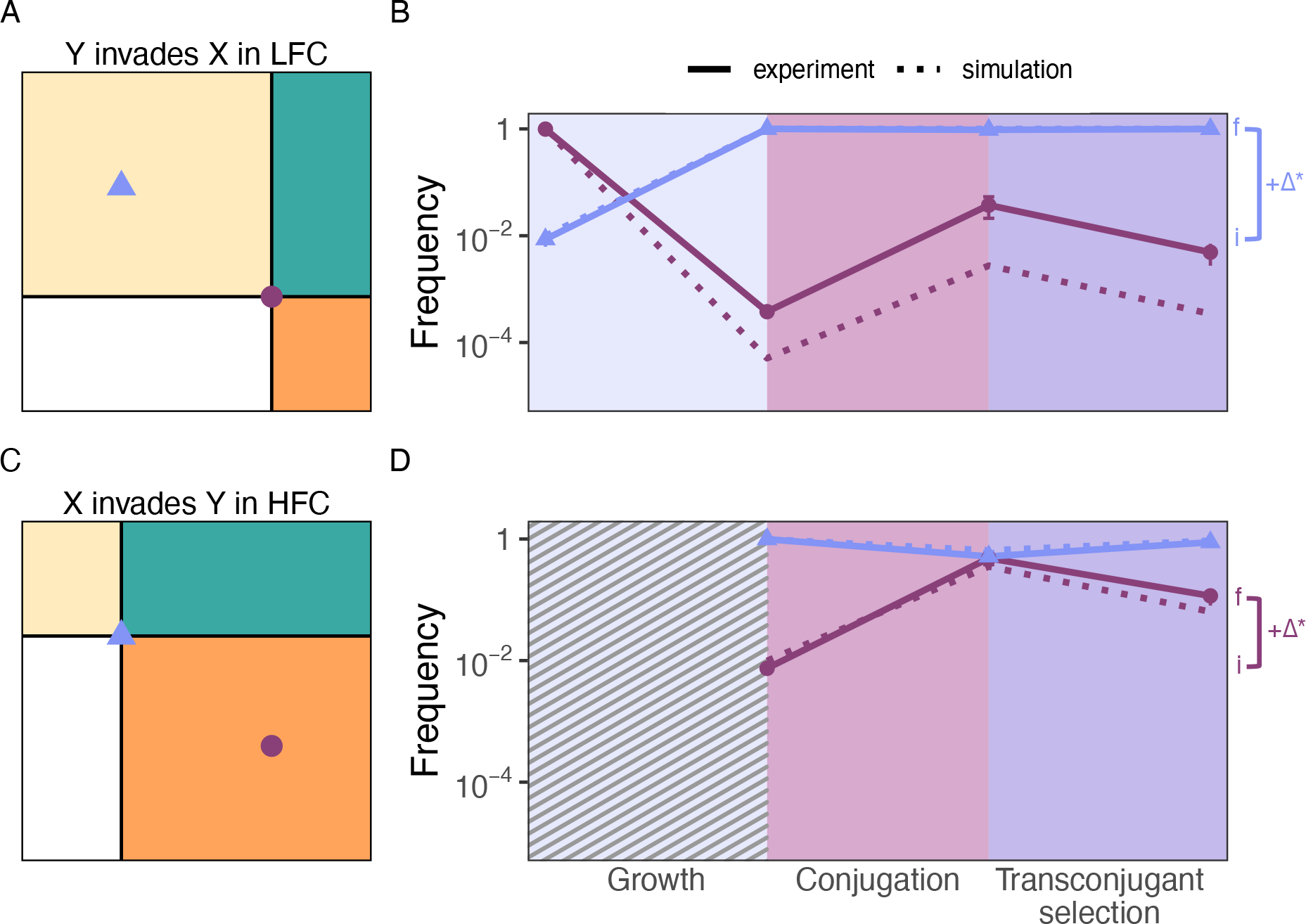
The antagonistic pleiotropy between genotypes X and Y allow us to explore transfer selection in various protocols. (A) In the LFC, the better grower genotype Y is expected to invade a resident population of genotype X due to the growth phase. (B) In both simulations (dotted lines) and experiments (solid lines), we see a significant increase in the frequency of genotype X over one cycle of the LFC (i, initial and f, final frequency). (C) In the HFC, the growth phase is removed (hashed) and thus the better conjugator genotype X is expected to invade a resident population of genotype Y. (D) Indeed, the frequency of genotype X significantly increases over one cycle of the HFC. Notably, genotype X increases in frequency during the conjugation phase in both protocols. Full simulations of multiple protocol cycles are shown in SI Figure S1. Error bars on experimental data points represent standard error of the mean.

We simulated and experimentally validated invasion dynamics over one cycle of each of these protocols: the strain predicted to be favored was started at low frequency in comparison to the opposing strain (1:100 ratio), and frequencies were tracked over each phase (see Materials and Methods for additional details). In both simulations and experiments, we found that genotype Y significantly increased in frequency over one cycle of the LFC (Figure 5B, *p* < 0.01, Welch’s t-test) and genotype X significantly increased in frequency over one cycle of the HFC (Figure 5D, p = 0.03, Welch’s t-test).

### Prediction of hypothetical mutant invasion across phenotypic space

Our case study focuses on a specific pair of strains, but our model can be used to perform a parameter sweep that predicts the invasion success of a wide range of hypothetical mutants relative to a specific ancestor across phenotypic space. We generated a set of hypothetical invading mutants with respect to the resident genotype X or Y and simulated their invasion across one cycle of the LFC or HFC.

Our parameter sweeps are consistent with results from our invasion simulations, confirming that genotype Y is favored in the LFC, invading genotype X (Figure 6A) and preventing the invasion of genotype X when it is the resident strain (Figure 6B). The opposite is true for genotype X in the HFC, invading genotype Y (Figure 6D) and preventing the invasion of genotype Y when it is the resident (Figure 6C).

**Figure 6.**
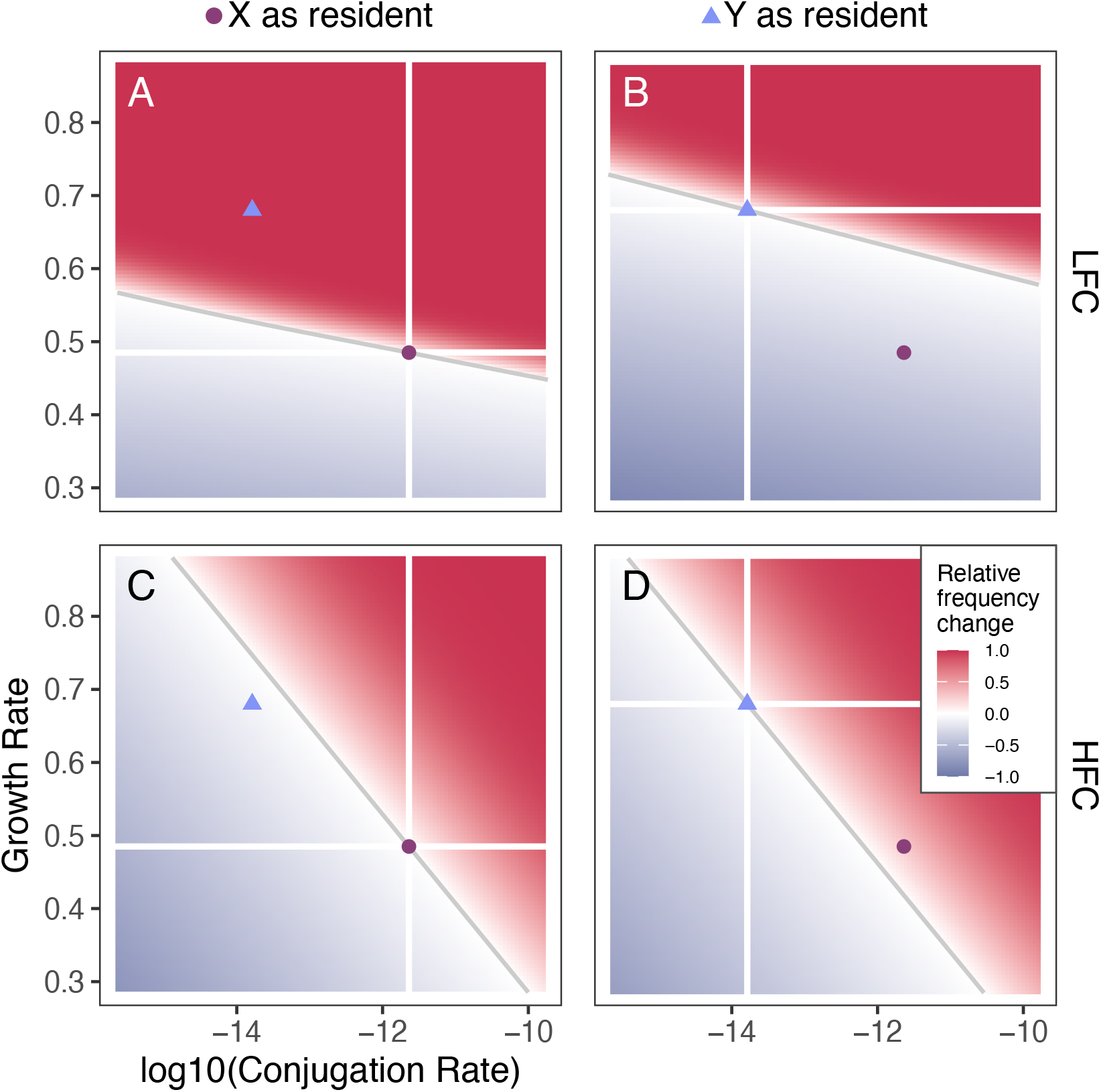
Parameter sweeps to predict mutant invasion across phenotypic space. With genotype X (A, C) or genotype Y (B, D) as the resident strain, we generated a set of hypothetical invaders and recorded invader frequency change over one cycle of the LFC (A, B) or HFC (C, D). Axes ranges were standardized across sweeps, and white axes were drawn at the conjugation and growth rate of the resident strain. Colors were scaled relative to the highest frequency change within each simulation, with red indicating a frequency increase, blue indicating a decrease, and more intense colors representing greater change. We note that the magnitude of the frequency change is not the same across the panels. The gray line represents the invasion boundary, separating space of exclusion (gray blue) and invasion (red).

We also found that the HFC favored a greater range of mutations in the parasitic quadrant (see Figure 1 for quadrant definitions), while the LFC favored a greater range of mutants in the host-centric quadrant (Figure 6). Both treatments favored mutants in the synergistic quadrant. These results mirror our findings from our abstract model, where increased availability of plasmid-free hosts increases the magnitude of the invasion boundary slope, allowing a larger proportion of the parasitic quadrant to be selected for. In these experimental treatments, this was achieved through increased frequency of conjugation and transconjugant phases, which additionally reduced growth phases.

### Application of invasion prediction framework to other studies

Beyond the strains from De Gelder et al., we applied our framework to another plasmid evolution experiment conducted by Dimitriu et al. (2021), which included frequent access to plasmid-free cells, but no forced conjugation [16]. The authors evolved the IncF plasmid R1 in *Escherichia coli* MG1655 in an experimental protocol with an “immigration” treatment: with each daily batch culture passage, a proportion of plasmid-free cells was added to the evolving plasmid-host pair. Thus, plasmid-free cells were always available for conjugation, but conjugation was not forced through transconjugant selection. The authors characterized a parasitic mutation in the R1 *copA* gene that increased conjugation rate while decreasing growth rate.

We adjusted our model to run Dimitriu et al.’s 90% immigration protocol, parameterized it with their measured conjugation and growth rates [16,21], and performed a parameter sweep with the wild type *E. coli* and R1 pair as the resident strain. Consistent with their results, our model predicts the *copA* mutation would be selected for (Figure 7). Interestingly, our model predicts that this immigration protocol selects for a wide range of both host-centric and parasitic mutations. The well-documented parasitic R1 *finO*^*-*^ mutation, which was also found by the authors, was predicted to increase at a greater frequency than the *copA* mutation (darker red). These results demonstrate a level of robustness to our framework, which can be used to predict the success of pleiotropic transfer mutations across qualitatively different experimental conditions.

**Figure 7.**
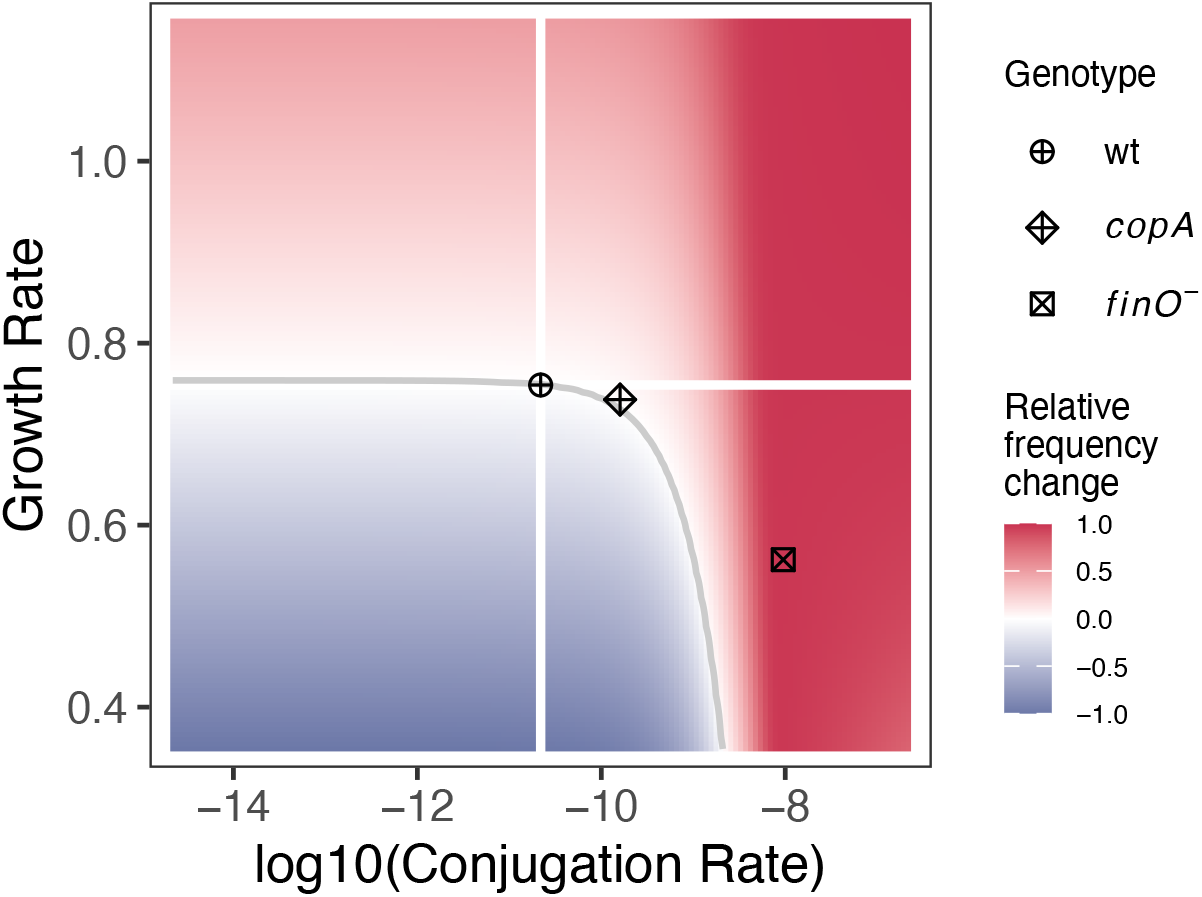
Parameter sweeps to predict mutant invasion in the Dimitriu et al. (2021) 90% immigration treatment. White axes were drawn at the conjugation and growth rate of the resident wild-type *E. coli*-R1 pair. Colors were scaled to the highest frequency change within each simulation, with red indicating a frequency increase and blue indicating a decrease. Other points represent R1 mutants that were found in the original experiment, with both the *copA* and *finO*^*-*^ mutants predicted to invade (land in red area).

## Discussion

### Standard experimental evolution protocols exert strong selective pressure on host growth, biasing plasmid evolution towards increasing VGT at the expense of HGT

Our results show that a long growth phase with multiple rounds of batch culture passages biases selection towards VGT, even when periodic access to plasmid-free cells is provided. Serial batch culture transfer is standard for microbial evolution experiments due to ease of execution (daily passages) and because it provides ample opportunity for mutation during growth [7,22]. Therefore, in evolution experiments with plasmid-bearing cells, this component results in heavy selective pressure on host growth, favoring host-centric (VGT-increasing, HGT-decreasing) mutations. Thus, even if parasitic (HGT-increasing, VGT-decreasing) mutations arose in an experiment, they would be filtered out by selection. This emphasis on VGT under standard experimental conditions can explain the bias towards observing host-centric evolution of plasmids in evolution experiments.

We show that a single change to a VGT-favoring protocol–eliminating the growth phase consisting of batch culture passages–can allow a greater range of parasitic mutations to invade in evolution experiments (Figure 6). In our multi-phase protocol, this also had the effect of increasing the frequency of exposure to plasmid-free cells, which prior work has identified as a factor increasing selection for HGT [6,16,23–25]. Consistent with our findings, Kottara et al. (2016) conducted an experiment with a similar “HFC” protocol, observing HGT-increasing evolutionary outcomes [18]. Although increasing the exposure rate to plasmid-free cells did lead to an enrichment of parasitic plasmid mutations in some cases, we reiterate that this factor alone did not guarantee such experimental outcomes [6,16,19].

### Beyond selective pressure, observing certain transfer mutations also depends on genetic accessibility

Another explanation for the lack of consistent HGT-increasing outcomes in plasmid evolution experiments beyond selective pressure is a lack of genetically accessible HGT-increasing mutations for a given ancestor. Even if a hypothetical mutant with a combination of transfer trait values could be selected for, there may be mutational constraints that prevent a genotype from existing. For example, our parameter sweeps emphasize the success of synergistic mutations in all treatments. However, single mutations that show synergy have only recently been discovered to occur, and it is currently unclear how prevalent and accessible synergistic mutations are for co-evolved plasmid-host pairs. These mutations may be more prevalent for new plasmid-host pairs where coevolution has yet to occur and are less likely to experience the constraints of a VGT-HGT tradeoff.

Antagonistic evolution may be more common than synergistic evolution, but it is unknown whether the distribution of available mutations is weighted towards host-centric or parasitic mutations. We note that growth continues to be important in batch culture protocols with frequent access to plasmid-free hosts. In the HFC, faster growers still benefit in conjugation and transconjugant phases, as a growth advantage can allow faster growers to access more plasmid-free cells for conjugation and for resulting transconjugants to dominate. We see this effect in our parameter sweeps–although the HFC favors a greater fraction of parasitic mutations (relative to the LFC), a reasonable portion of host-centric mutations can still invade (Figure 6). Thus, if host-centric mutations are more genetically accessible than parasitic mutations, they would be observed more often.

Finally, an accessible and selectively favored mutation arises from a single cell. Thus, it must increase in density to overcome bottlenecks imposed by density-dependent processes. In liquid systems, conjugation often follows a mass-action model, requiring a mutant to reach a specific density before interacting with recipients at an observable frequency [26]. Similarly, dilution during serial passage imposes bottlenecks [7]. Whether a mutation can overcome these bottlenecks can depend on timing of the mutation and potential clonal interference from competing mutations (SI Figure S2) [7,27]. The distribution of accessible mutations for a given ancestor, when the mutation is generated in an experiment, and the rate at which it can invade all contribute to our ability to observe mutations in phenotypic space. Determining the prevalence of each mutation type and evolution experiments under different selective regimes are ripe future directions for increasing our understanding of transfer rate evolution.

### Exploring more complex systems to expand the scope of experimental plasmid transfer evolution

Future studies should additionally focus on modeling and conducting experiments that incorporate more ecological and environmental factors to represent natural systems. Our study shows that a simple reduction in growth phases is enough to affect the transfer mutations that can be selected for. Thus, further consideration of plasmid biology, microbial community interactions, and environmental structure will continue to improve our ability to observe the types of transfer evolution occurring in nature.

Plasmids encode diverse maintenance mechanisms [28,29], thus, different plasmids may experience different selection under the same environmental conditions. For example, plasmid-encoded exclusion mechanisms hinder transfer into plasmid-bearing hosts [5,29]. Thus, many studies–including our own–make the basic assumption that only plasmid-free hosts can be recipients to conjugative transfer. However, exclusion mechanisms are imperfect and do not completely prevent superinfection [29]. This would open up the plasmid-bearing population as potential recipients, which could exert selection on HGT without exogenous addition of plasmid-free hosts [30]. Interestingly, a plasmid evolution experiment conducted by Yang et al. (2023) selected for a parasitic mutation despite their protocol consisting of batch culture passage in plasmid-selecting antibiotics (which would theoretically prevent plasmid-free cells from surviving) [31]. If superinfection was prominent in their system, that could potentially explain selection for the parasitic mutation. In systems where superinfection is observed at significant levels, models that incorporate superinfection, heteroplasmy, and segregational drift may be necessary for accurate predictions of plasmid transfer evolution [21,30,32].

The relationship between a plasmid and its host may also influence plasmid transfer evolution, especially in multispecies microbial communities. Plasmids can have different conjugation rates in different hosts, and a plasmid-host pair that results in a high conjugation rate can accelerate plasmid spread in microbial communities [33,34]. Especially for plasmids with a broad host range (e.g., IncP plasmids), being situated in a multispecies community could provide access to more recipients, driving selection for higher HGT rates [35]. Moving to a new host could also provide mutational opportunity to acquire HGT-increasing mutations not available in the prior host [36]. Considering plasmid-host and microbial community interactions can allow us to better predict transfer evolution.

Finally, environmental factors may influence transfer evolution, and considering factors like spatial structure could change whether VGT or HGT is favored. While laboratory experimental evolution is often conducted in well-mixed liquid systems, microbial communities harboring plasmids often exist in structured systems like biofilms [37,38]. Some plasmids even encode factors involved in biofilm formation [39]. In structured systems, conjugation may be more efficient–especially for plasmids that build shorter and more fragile pili [40,41]. In densely-packed biofilms, horizontal rather than vertical transfer may be a more feasible way for plasmids to spread, which could place higher selective pressure on HGT [42]. Indeed, there is evidence that such structure allows plasmids retaining HGT to persist where such horizontal functionality would otherwise be lost [12]. Additionally, structured systems allow the maintenance of a greater diversity of genotypes, which may increase the variety of evolutionary pathways available to a plasmid [38,43,44]. Incorporating spatial structure into evolution experiments will allow us to better predict transfer evolution, especially for clinically relevant plasmids (e.g., those carrying antibiotic resistance genes) that spread in biofilms [37,45].

Because conjugative plasmids are primary drivers of antibiotic resistance spread, understanding the precise conditions that improve horizontal gene transfer (HGT) is of critical medical importance [1]. The natural systems where this resistance spreads—such as in wastewater, agricultural, and hospital settings—possess unique features that differ fundamentally from laboratory conditions [45–47]. These environments may inherently promote HGT at higher rates than can be observed in experiments. Ultimately, bridging the gap between laboratory evolution experiments and complex natural ecosystems is essential to predicting how genes of major medical and environmental importance move.

## Acknowledgements

This work was supported by the Environmental Biology Division from the National Science Foundation (grant number 2142718, awarded to B.K. and E.M.T). O.K. was supported by the NSF Postdoctoral Research Fellowships in Biology Program grant no. DBI-2305907.

## Materials and Methods

### Bacterial Strains and Culture Conditions

Strains were obtained from De Gelder et al (2008) [17]. The authors evolved the 64.5 kb IncP-1β plasmid pB10 in the bacterial host *Stenotrophomonas maltophilia* P21 for ∼520 generations. To control for host evolution, the plasmid was conjugated between rifampicin (Rif) and nalidixic acid (Nal) resistant mutants of *S. maltophilia* P21. The evolved plasmids in ancestral hosts showed increased plasmid persistence compared to the ancestral plasmid-host pair. We focused on the evolved plasmid from replicate A, which was found to differ from the ancestral plasmid by a single base pair change in the *trbC* gene, yielding a single amino acid substitution in the encoded prepilin protein (A25E). We refer to the ancestral pair as genotype X (higher *x*-axis value) and the mutant pair as genotype Y (higher *y*-axis value).

All experiments were conducted in Lysogeny Broth (LB) and strains were grown at 30°C. Rif-marked strains were used as donors and Nal-marked strains as recipients in conjugation rate and invasion experiments. Donors, recipients, and transconjugants were differentiated with selective plating on LB agar plates. Donors were counted on LB plates supplemented with Rif 75 μg/mL and Tet 50 μg/mL, recipients on Nal 50 μg/mL, and transconjugants on Nal 50 μg/mL and Tet 50 μg/mL. Transconjugants would also be counted on recipient plates, but we note that the density of transconjugants was much lower than recipients (differing by over 4 orders of magnitude) and thus would not affect recipient density calculations. Genotype X and genotype Y were differentiated by colony size and subsequent streaking. Genotype X colonies were visibly smaller than Y colonies on dual antibiotic plates (due to lower host growth rates) and remained small colonies when re-streaked on new plates (Y maintained large colonies only). Additionally, we confirmed that this colony phenotype was representative of genotypes through PCR and restriction digests with BglI, which cuts genotype X but not Y at the site of the *trbC* mutation. Thus, we were able to track the densities of these different cell types without additional modifications to the plasmids.

### Conjugation Rate Assays

We measured the conjugation (HGT) rates of the plasmid-host pairs using the Luria-Delbruck Method [48,49]. For genotype X, overnight donor and recipient cultures were diluted to 10^7^ CFU/mL and incubated for 3 hours at 30°C to reach constant growth. The recipient was then diluted an additional 5-fold before mixing with the donor at a 1:1 ratio. Co-cultures incubated for 1 hr before transconjugant-selecting media was added. Genotype Y assays followed a similar protocol, except with an initial dilution to 5×10^7^ and a co-culture incubation time of 2 hr. Turbidity was recorded between 72-96 hours, when all negative controls showed no turbidity. The final units are expressed in conjugation events per donor density per hour (mL CFU^-1^ hr^-1^).

### Growth Rate Assays

We measured growth rate (VGT) of the plasmid-host pairs as well as plasmid-free *S. maltophilia* recipients over 4 hours to capture lag phase and maximum growth rate dynamics. Overnight cultures were diluted to 10^7^ CFU/mL and incubated for 4 hours at 30°C. Densities were measured through dilution plating at 0 and 4 hours. Assays were conducted at a volume of 5 mL and shaken at 200 rpm in 18 mm tubes. Growth rate (*ψ*) was calculated using the following formula:

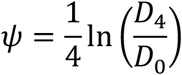

where *D*_*t*_ is the density of the focal strain at *t* hours. The final units are expressed in units per hour (hr^-1^).

### Invasion Experiments

We tested two selection protocols: the Low Frequency Conjugation (LFC) protocol and the High Frequency Conjugation protocol (HFC), predicted to favor HGT. The LFC was a modified version of the De Gelder et al. (2008) protocol to ensure experimental tractability of cell types, with a protocol cycle consisting of a growth phase (7 rounds of serial passage), conjugation phase, and transconjugant selection phase. We removed the growth phase to create the HFC, with a protocol cycle consisting of a conjugation and transconjugant selection phase.

The procedure for each phase is described as follows. Growth phases were conducted in 18 mm glass tubes at 5 mL, shaking at 200 rpm. This phase consisted of seven rounds of 1000-fold dilution into fresh media, followed by incubation for 24 hours. Conjugation phases were also conducted in glass tubes and started with a 100-fold batch culture dilution into fresh media and mixing with plasmid-free recipient cells at a 1:1 ratio. These tubes were not shaken, and conjugation phases ran for 5 hours. Transconjugant selection phases followed conjugation phases, with 500 μL of culture plated on transconjugant-selecting agar plates. Plates were incubated for 72 hours, until small and large colonies were visible. To estimate the final densities of cells after the transconjugant growth phase, representative small and large colonies were resuspended in saline and the total number of cells per colony was estimated. This estimated colony productivity was then multiplied by the number of relevant colonies from the transconjugant-selecting plates.

To confirm these protocols selected for the predicted mode of transfer, we initiated the plasmid genotype expected to invade at a ratio of 1:100, relative to the resident genotype. Specifically, we started the LFC with a population of 1:100 genotype Y to genotype X and the HFC with a population of 1:100 genotype X to genotype Y. We completed one cycle of each protocol, tracking densities over each phase. We compared frequencies of the invading type at the beginning and end of each cycle. We limited experimental validation to one cycle, as we noticed inconsistent colony size of genotype X after scraping and resuspending (mixed small and large), likely due to rapid evolution [10].

### Simulations

We used the R package “deSolve” to solve our differential equation system using the Euler method [50]. Simulations were run over 0.01 per hour time steps, recording densities and frequencies of each cell type.

To understand the success of the host-centric mutant (genotype Y) in De Gelder et al. (2008), we simulated its invasion in the conditions of the original evolution experiment. We programmed each “phase” into our simulations. In growth phases, the simulation replenished nutrients, diluted strains 1000-fold, and ran for 24 hours. The growth phase consisted of 7 rounds of this serial passage protocol. In conjugation phases, the simulation replenished nutrients, diluted strains 100-fold, added plasmid-free cells of the opposite cell type at a 1:1 ratio, and ran for 24 hours. In transconjugant selection phases, donor and recipient cell types were set to 0, transconjugants (opposite plasmid-containing cell types) were diluted 1000-fold, and ran for 24 hours. Simulations began with one mutant cell and ran for 3 protocol cycles.

LFC and HFC invasion simulations were run with the same starting densities as wet lab experiments (see Invasion Experiments). Conjugation phases were 5 hours long. Transconjugant selection phases also followed experimental protocol, taking transconjugants in 500 μL of culture and estimating transconjugant growth by multiplying the number of cells by the colony size of each strain. Full simulations were run until the resident strain was outcompeted (see Supplementary Figure S1).

Finally, we applied our model to predict the invasion success of hypothetical mutants that spanned our phenotypic axes in each experimental protocol. We set genotype X or Y as the resident strain, then generated a set of hypothetical mutants that spanned conjugation rates 4 orders of magnitude from the resident strain and growth rates 0.4 units from the resident strain. Points were generated at an increment of 150, resulting in 150*150 = 225,000 total simulations per plot. For each mutant, we simulated one cycle of each protocol and recorded the initial and final frequencies. We then plotted the magnitude of frequency change relative to the maximum change in each protocol on the phenotypic space.

### Statistical Analysis

We conducted two-tailed, paired (by day of experiment) Welch’s t-tests to confirm the conjugation and growth rates of genotypes X and Y were significantly different. For experimental validation of LFC and HFC invasion simulations, we maintained an *a priori* prediction from simulations that invading type would increase in frequency over one protocol cycle. Thus, we conducted one-tailed, paired (by replicate) Welch’s t-tests to confirm the frequency of the invading type increased over one protocol cycle.

