## Supplementary Information for "Molecular pet or parasite? Exploring selection for vertical and horizontal plasmid transfer"

**Supplemental Section 1: Rewriting differential equations in terms of proportions**

If we assume that mutations on our plasmid affect neither segregation nor death (i.e., $\sigma_{\mathrm{A}} = \sigma_{\mathrm{M}} = \sigma$ and $\delta_{\mathrm{A}} = \delta_{\mathrm{M}} = \delta$), we can rewrite equations [1] as follows:

$$\dot{A} = \psi_{\mathrm{A}}(1-\sigma)A + \gamma_{\mathrm{A}} aN - \delta A, \qquad [\mathrm{S1.1}a]$$

$$\dot{M} = \left(\psi_{\mathrm{A}} + \Delta_{\psi}\right)(1-\sigma)M + \left(\gamma_{\mathrm{A}} + \Delta_{\gamma}\right)mN - \delta M, \qquad [\mathrm{S1.1}b]$$

$$\dot{N} = \psi_{\mathrm{N}} N + \psi_{\mathrm{A}}\sigma A + \left(\psi_{\mathrm{A}} + \Delta_{\psi}\right)\sigma M - \gamma_{\mathrm{A}} aN - \left(\gamma_{\mathrm{A}} + \Delta_{\gamma}\right)mN - \delta_{\mathrm{N}} N. \qquad [\mathrm{S1.1}c]$$

Here we change our dynamic variables from densities to proportions. Let $x$ be the proportion of one of the strains and let $X$ be its density (such that $x = X/(A+M+N)$). By the quotient rule, we have:

$$\dot{x} = \frac{\dot{X}(A+M+N) - X(\dot{A}+\dot{M}+\dot{N})}{(A+M+N)^2} = \frac{\dot{X} - x(\dot{A}+\dot{M}+\dot{N})}{A+M+N}.$$

Because,

$$\frac{\dot{A}+\dot{M}+\dot{N}}{A+M+N} = (\psi_{\mathrm{A}} - \delta)a + \left(\psi_{\mathrm{A}} + \Delta_{\psi} - \delta\right)m + (\psi_{\mathrm{N}} - \delta_{\mathrm{N}})n\,,$$

we have

$$\dot{x} = \frac{\dot{X}}{A+M+N} - x\left[(\psi_{\mathrm{A}} - \delta)a + \left(\psi_{\mathrm{A}} + \Delta_{\psi} - \delta\right)m + (\psi_{\mathrm{N}} - \delta_{\mathrm{N}})n\right]. \qquad [\mathrm{S1.2}]$$

Using equations [S1.1] and equation [S1.2], we have:

$$\dot{a} = \psi_{\mathrm{A}}(1-\sigma)a + \gamma_{\mathrm{A}} an - \delta a - a\left[(\psi_{\mathrm{A}} - \delta)a + \left(\psi_{\mathrm{A}} + \Delta_{\psi} - \delta\right)m + (\psi_{\mathrm{N}} - \delta_{\mathrm{N}})n\right], \quad [\mathrm{S1.3}a]$$

$$\begin{aligned}\dot{m} = {} & \left(\psi_{\mathrm{A}} + \Delta_{\psi}\right)(1-\sigma)m + \left(\gamma_{\mathrm{A}} + \Delta_{\gamma}\right)mn - \delta m \\ & - m\left[(\psi_{\mathrm{A}} - \delta)a + \left(\psi_{\mathrm{A}} + \Delta_{\psi} - \delta\right)m + (\psi_{\mathrm{N}} - \delta_{\mathrm{N}})n\right], \qquad [\mathrm{S1.3}b]\end{aligned}$$

$$\begin{aligned}\dot{n} = {} & (\psi_{\mathrm{N}} - \delta_{\mathrm{N}})n + \psi_{\mathrm{A}}\sigma a + \left(\psi_{\mathrm{A}} + \Delta_{\psi}\right)\sigma m - \gamma_{\mathrm{A}} an - \left(\gamma_{\mathrm{A}} + \Delta_{\gamma}\right)mn \\ & - n\left[(\psi_{\mathrm{A}} - \delta)a + \left(\psi_{\mathrm{A}} + \Delta_{\psi} - \delta\right)m + (\psi_{\mathrm{N}} - \delta_{\mathrm{N}})n\right]. \qquad [\mathrm{S1.3}c]\end{aligned}$$

Equations [S1.3] recast the dynamics in terms of proportions of the three bacterial strains.

### Supplemental Section 2: Two-strain equilibrium and its stability

Consider a community without mutant plasmids present ($m = 0$). Equations [S1.3] (or, equivalently, equations [2]) reduce to the following:

$$\dot{a} = \psi_A(1-\sigma)a + \gamma_A an - \delta a - a[(\psi_A - \delta)a + (\psi_N - \delta_N)n], \qquad [S2.1a]$$

$$\dot{n} = (\psi_N - \delta_N)n + \psi_A\sigma a - \gamma_A an - n[(\psi_A - \delta)a + (\psi_N - \delta_N)n]. \qquad [S2.1b]$$

One equilibrium is $\hat{a} = 0$ and $\hat{n} = 1$ (i.e., $\dot{a} = 0$ and $\dot{n} = 0$ in equations [S2.1] when $a = 0$ and $n = 1$). However, because we are interested in plasmid evolution, we are specifically interested in equilibria where plasmid-bearing cells are present (i.e., $\hat{a} > 0$). At any equilibrium, $\dot{a} = 0$, such that equilibria will satisfy:

$$0 = \psi_A(1-\sigma)\hat{a} + \gamma_A\hat{a}\hat{n} - \delta\hat{a} - \hat{a}[(\psi_A - \delta)\hat{a} + (\psi_N - \delta_N)\hat{n}]$$

Because we explicitly are focusing on cases where $\hat{a} \neq 0$ (and note $\hat{n} = 1 - \hat{a}$), we have

$$0 = \psi_A(1-\sigma) + \gamma_A(1-\hat{a}) - \delta - (\psi_A - \delta)\hat{a} - (\psi_N - \delta_N)(1-\hat{a})$$

$$(\psi_A - \delta + \gamma_A - \psi_N + \delta_N)\hat{a} = \psi_A(1-\sigma) - \delta + \gamma_A - \psi_N + \delta_N$$

$$\hat{a} = \frac{\psi_A(1-\sigma) - \delta + \gamma_A - \psi_N + \delta_N}{\psi_A - \delta + \gamma_A - \psi_N + \delta_N} \qquad [S2.2]$$

As long as $\psi_A(1-\sigma) - \delta + \gamma_A > \psi_N - \delta_N$, the numerator of [S2.2] is positive, and if $\sigma > 0$, then

$$\{\psi_A - \delta + \gamma_A - \psi_N + \delta_N\} > \{\psi_A(1-\sigma) - \delta + \gamma_A - \psi_N + \delta_N\} > 0$$

and the denominator of [S2.2] is positive as well, and $\hat{a}$ is well defined ($0 < \hat{a} < 1$). We can rewrite this equilibrium value as:

$$\hat{a} = 1 - \frac{\psi_A\sigma}{\psi_A - \delta + \gamma_A - \psi_N + \delta_N}.$$

And because $\hat{n} = 1 - \hat{a}$,

$$\hat{n} = \frac{\psi_A\sigma}{\psi_A - \delta + \gamma_A - \psi_N + \delta_N}. \qquad [S2.3]$$

We now turn to the stability of this internal equilibrium. Without mutant plasmids present, we only need to track the proportion of ancestral plasmid-bearing cells (as the proportion of plasmid-free cells is immediately given once the proportion of ancestral plasmid-bearing cells is known). That is, we have a single relevant differential equation:

$$\dot{a} = h(a),$$

where, from equation [S2.1a],

$$h(a) = \psi_{\mathrm{A}}(1-\sigma)a + \gamma_{\mathrm{A}}a(1-a) - \delta a - a[(\psi_{\mathrm{A}} - \delta)a + (\psi_{\mathrm{N}} - \delta_{\mathrm{N}})(1-a)] \quad \text{[S2.4]}$$

Simplifying equation [S2.4] gives:

$$h(a) = \psi_{\mathrm{A}}(1-\sigma)a + \gamma_{\mathrm{A}}a - \gamma_{\mathrm{A}}a^2 - \delta a - (\psi_{\mathrm{A}} - \delta)a^2 - (\psi_{\mathrm{N}} - \delta_{\mathrm{N}})a + (\psi_{\mathrm{N}} - \delta_{\mathrm{N}})a^2$$

$$h(a) = \{\psi_{\mathrm{A}}(1-\sigma) - \delta + \gamma_{\mathrm{A}} - \psi_{\mathrm{N}} + \delta_{\mathrm{N}}\}a - \{\psi_{\mathrm{A}} - \delta + \gamma_{\mathrm{A}} - \psi_{\mathrm{N}} + \delta_{\mathrm{N}}\}a^2$$

Taking the derivative of $h$ with respect to $a$ gives:'

$$\frac{dh}{da} = \{\psi_{\mathrm{A}}(1-\sigma) - \delta + \gamma_{\mathrm{A}} - \psi_{\mathrm{N}} + \delta_{\mathrm{N}}\} - 2\{\psi_{\mathrm{A}} - \delta + \gamma_{\mathrm{A}} - \psi_{\mathrm{N}} + \delta_{\mathrm{N}}\}a$$

Evaluating the derivative at the equilibrium (equation [S2.2]) gives:

$$\left.\frac{dh}{da}\right|_{a=\hat{a}} = \{\psi_{\mathrm{A}}(1-\sigma) - \delta + \gamma_{\mathrm{A}} - \psi_{\mathrm{N}} + \delta_{\mathrm{N}}\} - 2\{\psi_{\mathrm{A}}(1-\sigma) - \delta + \gamma_{\mathrm{A}} - \psi_{\mathrm{N}} + \delta_{\mathrm{N}}\}$$

$$\left.\frac{dh}{da}\right|_{a=\hat{a}} = -\{\psi_{\mathrm{A}}(1-\sigma) - \delta + \gamma_{\mathrm{A}} - \psi_{\mathrm{N}} + \delta_{\mathrm{N}}\}$$

Because we must have $\psi_{\mathrm{A}}(1-\sigma) - \delta + \gamma_{\mathrm{A}} - \psi_{\mathrm{N}} + \delta_{\mathrm{N}} > 0$ (in order for the equilibrium [S2.2] to be well-defined), it follows that

$$\left.\frac{dh}{da}\right|_{a=\hat{a}} < 0,$$

which means the equilibrium in a population consisting of only ancestral plasmid-bearing cells and plasmid-free cells.

$$\left(\hat{a} = 1 - \frac{\psi_{\mathrm{A}}\sigma}{\psi_{\mathrm{A}} - \delta + \gamma_{\mathrm{A}} - \psi_{\mathrm{N}} + \delta_{\mathrm{N}}}, \qquad \hat{n} = \frac{\psi_{\mathrm{A}}\sigma}{\psi_{\mathrm{A}} - \delta + \gamma_{\mathrm{A}} - \psi_{\mathrm{N}} + \delta_{\mathrm{N}}}\right),$$

is stable.

**Supplemental Section 3: Local linear stability analysis for a rare mutant**

What happens when we consider perturbations that introduce a small proportion of mutant plasmid-bearing cells? Or, put another way, is the equilibrium

$$\left(\hat{a} = 1 - \frac{\psi_\mathrm{A}\sigma}{\psi_\mathrm{A} - \delta + \gamma_\mathrm{A} - \psi_\mathrm{N} + \delta_\mathrm{N}}, \qquad \hat{m} = 0, \qquad \hat{n} = \frac{\psi_\mathrm{A}\sigma}{\psi_\mathrm{A} - \delta + \gamma_\mathrm{A} - \psi_\mathrm{N} + \delta_\mathrm{N}}\right),$$

stable?

Here, we can represent the dynamical system as two coupled differential equations:

$$\dot{a} = f(a, m),$$

$$\dot{m} = g(a, m),$$

where, using equations [S1.3a] and [S1.3b], we have

$$\begin{aligned} f(a,m) = {} & \psi_\mathrm{A}(1-\sigma)a + \gamma_\mathrm{A}a(1-a-m) - \delta a \\ & - a\left[(\psi_\mathrm{A} - \delta)a + \left(\psi_\mathrm{A} + \Delta_\psi - \delta\right)m + (\psi_\mathrm{N} - \delta_\mathrm{N})(1-a-m)\right], \end{aligned}$$

$$\begin{aligned} g(a,m) = {} & \left(\psi_\mathrm{A} + \Delta_\psi\right)(1-\sigma)m + \left(\gamma_\mathrm{A} + \Delta_\gamma\right)m(1-a-m) - \delta m \\ & - m\left[(\psi_\mathrm{A} - \delta)a + \left(\psi_\mathrm{A} + \Delta_\psi - \delta\right)m + (\psi_\mathrm{N} - \delta_\mathrm{N})(1-a-m)\right] \end{aligned}$$

Simplifying, we have

$$\begin{aligned} f(a,m) = {} & \{\psi_\mathrm{A}(1-\sigma) - \delta + \gamma_\mathrm{A} - \psi_\mathrm{N} + \delta_\mathrm{N}\}a - \{\psi_\mathrm{A} - \delta + \gamma_\mathrm{A} - \psi_\mathrm{N} + \delta_\mathrm{N}\}a^2 \\ & - \left\{\psi_\mathrm{A} + \Delta_\psi - \delta + \gamma_\mathrm{A} - \psi_\mathrm{N} + \delta_\mathrm{N}\right\}am, \end{aligned}$$

$$\begin{aligned} g(a,m) = {} & \left\{\left(\psi_\mathrm{A} + \Delta_\psi\right)(1-\sigma) - \delta + \gamma_\mathrm{A} + \Delta_\gamma - \psi_\mathrm{N} + \delta_\mathrm{N}\right\}m \\ & - \left\{\psi_\mathrm{A} + \Delta_\psi - \delta + \gamma_\mathrm{A} + \Delta_\gamma - \psi_\mathrm{N} + \delta_\mathrm{N}\right\}m^2 \\ & - \left\{\psi_\mathrm{A} - \delta + \gamma_\mathrm{A} + \Delta_\gamma - \psi_\mathrm{N} + \delta_\mathrm{N}\right\}am \end{aligned}$$

The Jacobian matrix for this system is

$$\mathbf{J} = \begin{bmatrix} \left.\frac{\partial f}{\partial a}\right|_{\hat{a},\hat{m}} & \left.\frac{\partial f}{\partial m}\right|_{\hat{a},\hat{m}} \\ \left.\frac{\partial g}{\partial a}\right|_{\hat{a},\hat{m}} & \left.\frac{\partial g}{\partial m}\right|_{\hat{a},\hat{m}} \end{bmatrix}.$$

We'll take each entry in turn, starting with $\left.\frac{\partial f}{\partial a}\right|_{\hat{a},\hat{m}}$:

$$\begin{aligned} \left.\frac{\partial f}{\partial a}\right|_{\hat{a},\hat{m}} = {} & \{\psi_\mathrm{A}(1-\sigma) - \delta + \gamma_\mathrm{A} - \psi_\mathrm{N} + \delta_\mathrm{N}\} - 2\{\psi_\mathrm{A} - \delta + \gamma_\mathrm{A} - \psi_\mathrm{N} + \delta_\mathrm{N}\}\hat{a} \\ & - \left\{\psi_\mathrm{A} + \Delta_\psi - \delta + \gamma_\mathrm{A} - \psi_\mathrm{N} + \delta_\mathrm{N}\right\}\hat{m}, \end{aligned}$$

Using equation [S2.2] and setting $\hat{m} = 0$, we have the following:

$$\left.\frac{\partial f}{\partial a}\right|_{\hat{a},\hat{m}} = \{\psi_\mathrm{A}(1-\sigma) - \delta + \gamma_\mathrm{A} - \psi_\mathrm{N} + \delta_\mathrm{N}\} - 2\{\psi_\mathrm{A}(1-\sigma) - \delta + \gamma_\mathrm{A} - \psi_\mathrm{N} + \delta_\mathrm{N}\}$$

$$\left.\frac{\partial f}{\partial a}\right|_{\hat{a},\hat{m}} = -\{\psi_\mathrm{A}(1-\sigma) - \delta + \gamma_\mathrm{A} - \psi_\mathrm{N} + \delta_\mathrm{N}\}$$

Turning to $\left.\frac{\partial f}{\partial m}\right|_{\hat{a},\hat{m}}$:

$$\left.\frac{\partial f}{\partial m}\right|_{\hat{a},\hat{m}} = -\{\psi_\mathrm{A} + \Delta_\psi - \delta + \gamma_\mathrm{A} - \psi_\mathrm{N} + \delta_\mathrm{N}\}\hat{a}$$

Using equation [S2.2],

$$\left.\frac{\partial f}{\partial m}\right|_{\hat{a},\hat{m}} = -\{\psi_\mathrm{A} + \Delta_\psi - \delta + \gamma_\mathrm{A} - \psi_\mathrm{N} + \delta_\mathrm{N}\}\frac{\psi_\mathrm{A}(1-\sigma) - \delta + \gamma_\mathrm{A} - \psi_\mathrm{N} + \delta_\mathrm{N}}{\psi_\mathrm{A} - \delta + \gamma_\mathrm{A} - \psi_\mathrm{N} + \delta_\mathrm{N}}$$

$$\left.\frac{\partial f}{\partial m}\right|_{\hat{a},\hat{m}} = -\{\psi_\mathrm{A} - \delta + \gamma_\mathrm{A} - \psi_\mathrm{N} + \delta_\mathrm{N} + \Delta_\psi\}\frac{\psi_\mathrm{A}(1-\sigma) - \delta + \gamma_\mathrm{A} - \psi_\mathrm{N} + \delta_\mathrm{N}}{\psi_\mathrm{A} - \delta + \gamma_\mathrm{A} - \psi_\mathrm{N} + \delta_\mathrm{N}}$$

$$\left.\frac{\partial f}{\partial m}\right|_{\hat{a},\hat{m}} = -\{\psi_\mathrm{A}(1-\sigma) - \delta + \gamma_\mathrm{A} - \psi_\mathrm{N} + \delta_\mathrm{N}\} - \Delta_\psi\frac{\psi_\mathrm{A}(1-\sigma) - \delta + \gamma_\mathrm{A} - \psi_\mathrm{N} + \delta_\mathrm{N}}{\psi_\mathrm{A} - \delta + \gamma_\mathrm{A} - \psi_\mathrm{N} + \delta_\mathrm{N}}$$

$$\left.\frac{\partial f}{\partial m}\right|_{\hat{a},\hat{m}} = -\{\psi_\mathrm{A}(1-\sigma) - \delta + \gamma_\mathrm{A} - \psi_\mathrm{N} + \delta_\mathrm{N}\} - \Delta_\psi\hat{a}$$

Next, we look at $\left.\frac{\partial g}{\partial a}\right|_{\hat{a},\hat{m}}$:

$$\left.\frac{\partial g}{\partial a}\right|_{\hat{a},\hat{m}} = -\{\psi_\mathrm{A} - \delta + \gamma_\mathrm{A} + \Delta_\gamma - \psi_\mathrm{N} + \delta_\mathrm{N}\}\hat{m}$$

As $\hat{m} = 0$, we have

$$\left.\frac{\partial g}{\partial a}\right|_{\hat{a},\hat{m}} = 0.$$

Finally, we consider $\left.\frac{\partial g}{\partial m}\right|_{\hat{a},\hat{m}}$:

$$\left.\frac{\partial g}{\partial m}\right|_{\hat{a},\hat{m}} = \{(\psi_\mathrm{A} + \Delta_\psi)(1-\sigma) - \delta + \gamma_\mathrm{A} + \Delta_\gamma - \psi_\mathrm{N} + \delta_\mathrm{N}\} \\ - 2\{\psi_\mathrm{A} + \Delta_\psi - \delta + \gamma_\mathrm{A} + \Delta_\gamma - \psi_\mathrm{N} + \delta_\mathrm{N}\}\hat{m} - \{\psi_\mathrm{A} - \delta + \gamma_\mathrm{A} + \Delta_\gamma - \psi_\mathrm{N} + \delta_\mathrm{N}\}\hat{a}$$

Using equation [S2.2] and setting $\hat{m} = 0$, we have

$$\left.\frac{\partial g}{\partial m}\right|_{\hat{a},\hat{m}} = \{(\psi_{\mathrm{A}} + \Delta_{\psi})(1-\sigma) - \delta + \gamma_{\mathrm{A}} + \Delta_{\gamma} - \psi_{\mathrm{N}} + \delta_{\mathrm{N}}\} - \{\psi_{\mathrm{A}} - \delta + \gamma_{\mathrm{A}} - \psi_{\mathrm{N}} + \delta_{\mathrm{N}} + \Delta_{\gamma}\}\frac{\psi_{\mathrm{A}}(1-\sigma) - \delta + \gamma_{\mathrm{A}} - \psi_{\mathrm{N}} + \delta_{\mathrm{N}}}{\psi_{\mathrm{A}} - \delta + \gamma_{\mathrm{A}} - \psi_{\mathrm{N}} + \delta_{\mathrm{N}}}.$$

Simplifying gives:

$$\left.\frac{\partial g}{\partial m}\right|_{\hat{a},\hat{m}} = \{\psi_{\mathrm{A}}(1-\sigma) - \delta + \gamma_{\mathrm{A}} - \psi_{\mathrm{N}} + \delta_{\mathrm{N}} + (1-\sigma)\Delta_{\psi} + \Delta_{\gamma}\} - \{\psi_{\mathrm{A}} - \delta + \gamma_{\mathrm{A}} - \psi_{\mathrm{N}} + \delta_{\mathrm{N}} + \Delta_{\gamma}\}\left(1 - \frac{\psi_{\mathrm{A}}\sigma}{\psi_{\mathrm{A}} - \delta + \gamma_{\mathrm{A}} - \psi_{\mathrm{N}} + \delta_{\mathrm{N}}}\right)$$

$$\left.\frac{\partial g}{\partial m}\right|_{\hat{a},\hat{m}} = \{\psi_{\mathrm{A}}(1-\sigma) - \delta + \gamma_{\mathrm{A}} - \psi_{\mathrm{N}} + \delta_{\mathrm{N}} + (1-\sigma)\Delta_{\psi} + \Delta_{\gamma}\} - \{\psi_{\mathrm{A}} - \delta + \gamma_{\mathrm{A}} - \psi_{\mathrm{N}} + \delta_{\mathrm{N}} + \Delta_{\gamma}\} + \psi_{\mathrm{A}}\sigma + \left(\frac{\psi_{\mathrm{A}}\sigma}{\psi_{\mathrm{A}} - \delta + \gamma_{\mathrm{A}} - \psi_{\mathrm{N}} + \delta_{\mathrm{N}}}\right)\Delta_{\gamma}$$

$$\left.\frac{\partial g}{\partial m}\right|_{\hat{a},\hat{m}} = (1-\sigma)\Delta_{\psi} + \hat{n}\Delta_{\gamma}$$

Thus, our Jacobian matrix is:

$$\mathbf{J} = \begin{bmatrix} -\{\psi_{\mathrm{A}}(1-\sigma) - \delta + \gamma_{\mathrm{A}} - \psi_{\mathrm{N}} + \delta_{\mathrm{N}}\} & -\{\psi_{\mathrm{A}}(1-\sigma) - \delta + \gamma_{\mathrm{A}} - \psi_{\mathrm{N}} + \delta_{\mathrm{N}}\} - \Delta_{\psi}\hat{a} \\ 0 & (1-\sigma)\Delta_{\psi} + \hat{n}\Delta_{\gamma} \end{bmatrix}.$$

Because this matrix is triangular, the eigenvalues line the diagonal:

$$\lambda_1 = -\{\psi_{\mathrm{A}}(1-\sigma) - \delta + \gamma_{\mathrm{A}} - \psi_{\mathrm{N}} + \delta_{\mathrm{N}}\},$$

$$\lambda_2 = (1-\sigma)\Delta_{\psi} + \hat{n}\Delta_{\gamma}.$$

As discussed in Supplemental Section 2, the numerator of [S2.2] will need to be positive for the existence of a non-trivial equilibrium, ensuring the first eigenvalue is negative ($\lambda_1 < 0$). The second eigenvalue relates to the stability of the equilibrium to perturbations (i.e., small non-zero introductions of) mutant plasmid-bearing cells. Specifically, this equilibrium is unstable when $\lambda_2 > 0$, or

$$(1-\sigma)\Delta_{\psi} + \hat{n}\Delta_{\gamma} > 0. \qquad \text{[S3.1]}$$

This gives the following simple mutant invasion criterion:

$$\Delta_{\psi} > -\frac{\hat{n}}{1-\sigma}\Delta_{\gamma}, \qquad \text{[S3.2]}$$

which is condition [4] in our main text.

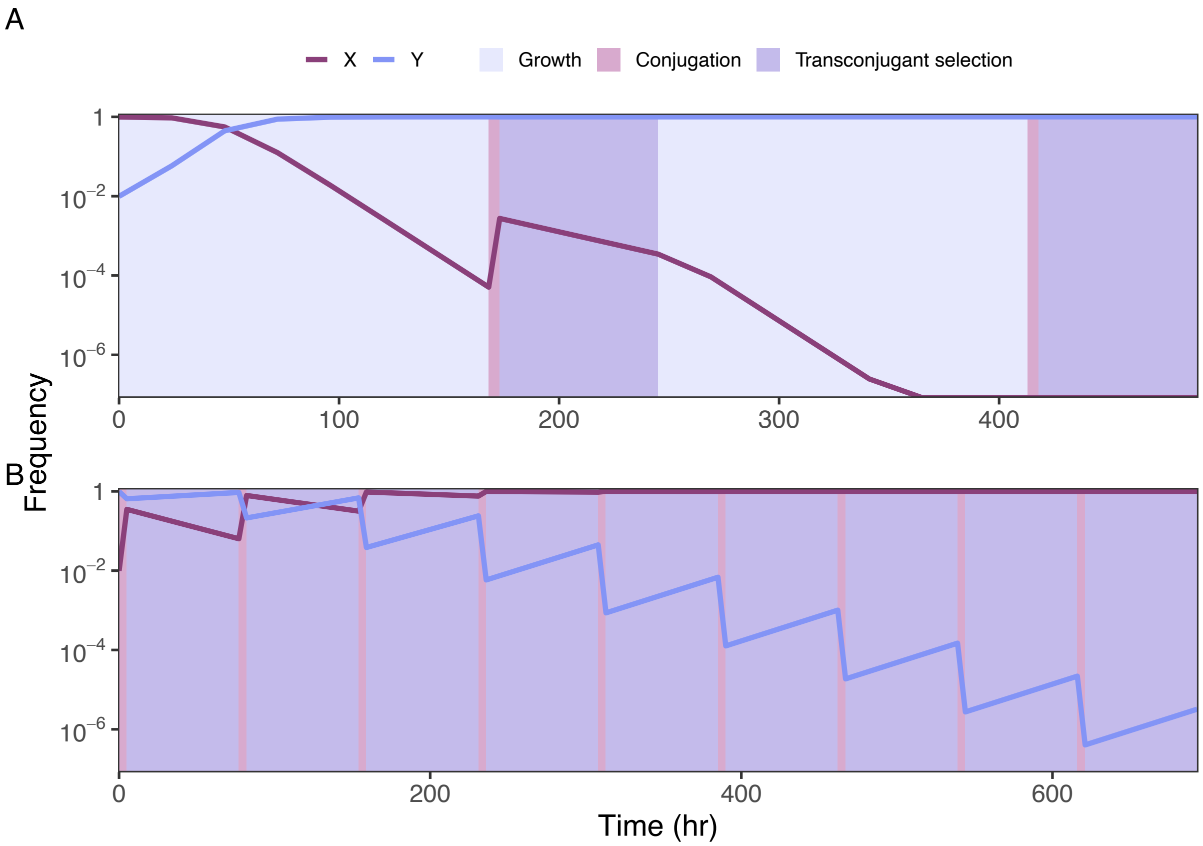


**Figure S1.** Full invasion experiment simulations, from start to displacement of the resident strain. Simulations begin with the invading type at a 1:100 ratio compared to the resident type. Frequencies of each genotype are plotted over time. (A) The host-centric genotype Y outcompetes and displaces genotype X over 2 cycles of the LFC. (B) The parasitic genotype X outcompetes and displaces genotype Y over 10 cycles of the HFC. Experimental validation of the first cycle of each protocol was completed (see Figure 5 in the main text).

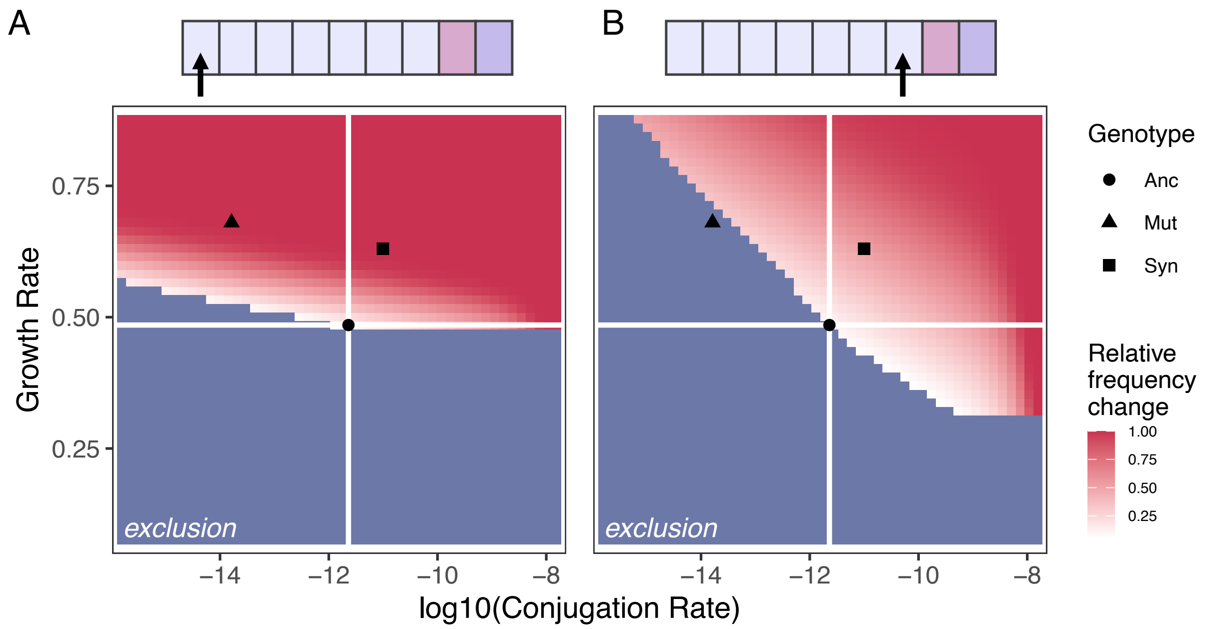


**Figure S2.** Timing and clonal interference can influence what mutations are observable. In De Gelder et al. (2008), the ancestral strain (circle, center of axes) was evolved in an LFC protocol. A host-centric (triangle) mutation and a synergistic (square) mutation were identified at the end of the experiment. The successes of these mutations could have depended on when they arose in the experiment. Blue indicates hypothetical mutants that were excluded and red indicates mutants that invaded, with darker red indicating greater frequency increases. (A) As both strains have a growth advantage, emerging early in the growth phase would have favored both strains (landing in the red area). (B) However, emerging later in the growth phase would limit the advantage of the host-centric strain, only allowing the synergistic strain to invade. Additionally, if both mutations emerged in the same population, the synergistic strain would outcompete the host-centric strain due to its conjugation advantage, indicated by the synergistic mutant landing in a darker red area. Consequently, the synergistic mutation was observed in more replicates than the host-centric mutation (De Gelder et al. 2008). Stochastic timing and clonal interference can influence our ability to observe mutations, even if they are selected for.
